# Disentangling the CHAOS of intrinsic disorder in human proteins

**DOI:** 10.1101/2024.10.26.620428

**Authors:** Ida de Vries, Jitske Bak, Daniel Álvarez Salmoral, Ren Xie, Razvan Borza, Maria Konijnenberg, Anastassis Perrakis

## Abstract

Most proteins consist of both folded domains and Intrinsically Disordered Regions (IDRs). However, the widespread occurrence of intrinsic disorder in human proteins, along with its characteristics, is often overlooked by the broader communities of structural and molecular biologists. Building on the MobiDB database of intrinsic disorder in proteins, here we develop a comprehensive dataset (**C**omprehensive analysis of Human proteins **A**nd their dis**O**rdered Segments - CHAOS). We implement internally consistent definitions of disordered regions, and annotate general characteristics such as cellular location, essentiality, post-translational modifications, and predicted pathogenicity. Further, we cross-reference to structure predictions from AlphaFold. We find that most human proteins contain at least one disordered region, predominantly located at the protein termini. IDRs are less hydrophobic, enriched in post-translational modifications, and mutations in IDRs are predicted to be less pathogenic than in non-IDRs. Additionally, we discovered that proteins residing in different cellular locations possess distinct disorder profiles. Finally, the predicted AlphaFold models of proteins in CHAOS suggest that disordered regions and proteins are often predicted to adopt secondary structure. Hereby we enhance the visibility and understanding of intrinsic disorder in human proteins.

**Key messages:**

1. Four out of five human proteins contain one or more intrinsically disordered regions (IDRs).
2. Half of the IDRs are located at protein termini, but three quarters of all human proteins contain a terminal IDR.
3. The amount and location of disordered regions differs throughout cellular compartments.
4. One in five missense mutations in IDRs are likely pathogenic.
5. AlphaFold predicts secondary structure elements within intrinsically disordered regions and fully disordered proteins.

## Introduction

Proteins typically consist of folded domains and intrinsically disordered regions (IDRs). Folded domains are well characterized and contain a plethora of functions. Although the roles of IDRs are less well understood, there is increasing evidence that IDRs have important functions, like protein-protein interactions (also referred to as fuzzy complexes^1^ or flexible nets^2^), aid rapid evolution^3^, and can be involved in disease^4^. IDRs have also been implicated in numerous roles in cellular processes, such as signaling^5,6^, protein degradation^7^, translation initiation^8^, membrane remodeling^9^, viral infection^10^, and chromatin remodeling^11^.

Studies focusing on IDRs typically rely on fluorescent microscopy. While these are very useful to study the spatiotemporal nature of interactions, these techniques cannot visualize proteins at the atomic level. Structural biology is well equipped to study the atomic level of folded proteins. However, the techniques are poorly equipped to visualize IDRs and their interactions in vivo or in a native environment. X-ray analysis^12,13^ requires the crystallization of protein, which is hindered by the presence of IDRs. Cryo-EM^14,15^ tolerates the presence of IDRs, however these most often stay invisible or blurry due to a lower signal to noise ratio. NMR^16,17^ is the most suitable technique for detecting more flexible regions, but protein structure determination is hindered by protein size, labeling and solubility. These experimental limitations made scientists massively ignore the existence of disordered regions in proteins^18,19^.

The growing capabilities of computational studies have enabled the identification and characterization of intrinsically disordered regions (IDRs). Previously, there has been a focus on predicting protein regions that are disordered based on the amino acid sequence of the protein. Examples of deployed methods and databases are the IUPRED^20^, GLOBPLOT^18^ and DISEMBL^19^. Amusingly, these servers have mostly been developed to aid protein crystallization^19,21^, neglecting the potential of studying IDRs themselves. More recently, Artificial Intelligence (AI) techniques made it possible to predict relatively accurate atomic structures of proteins by algorithms like AlphaFold2^22^, AlphaFold3^23^ and RosettaFold^24^. Although these techniques were also developed to predict the structured elements of proteins, they drastically increased the visibility of protein disorder at atomic level^25,26^.

Taking advantage of computational algorithms and available experimental data, efforts have been made to improve annotations of IDRs. A notable information-rich resource among them is MobiDB^27,28^. MobiDB contains annotations of IDRs that are based on manual curation, experimental structure models and their homologs, and IDR predictions. Furthermore, this database contains annotations of e.g. post translational modifications (PTMs), as well as annotations of the ability to phase separate.

In this work, we obtain IDRs from MobiDB^28^, and add annotations on essentiality (from the “essentialome”^29,30^), cellular location (from UniProt^31^), and additional PTMs (from dbPTM^32^) to create a comprehensive dataset of IDRs in human proteins. We look at the abundance and properties of IDRs in the protein sequence and in the context of the human cell. We study IDRs in the context of predicted protein structure models and characterize fully disordered proteins.

## Results and discussion

### What are IDRs?

A problem studying IDRs is that there is no formalized definition. There is no consensus about the minimum number of consecutive disordered residues that constitute an IDR. As such, some studies define IDRs as being disordered regions longer than 5 consecutive residues^33^, others select and analyze only disordered regions longer than 10 or 30 residues^26,34^. In this work, we set the minimum length for IDRs at 8 residues. The average length of unmodeled loops (regions shorter than 30 residues) in the PDB-REDO databank^35^ is 7 amino acids (median 5 residues). Therefore, as IDRs are supposed to be longer than loops, defining an IDR as 8 or more consecutive disordered residues is a reasonable choice for the purpose of this study.

Using the above definition, we build our dataset, CHAOS: **C**omprehensive analysis of **H**uman proteins **A**nd their dis**O**rdered **S**egments. We mined the IDR locations of human proteins as annotated in MobiDB, which is our foundation dataset. Additionally, we downloaded the predicted protein structure models of all human proteins from the AlphaFold predicted protein structure database^22,36^. These models contain a per residue pLDDT (Predicted Local Distance Difference Test) score, which typically is low (not confident) for IDRs and high (confident) for structured regions (non-IDRs)^25,26^. To annotate secondary structure elements in these protein models, we ran DSSP (Dictionary of Secondary Structure in Protein) to determine if residues in the models adopt a secondary structure or not.

As not all IDRs obtained from MobiDB obey our above set definition (8 or more consecutive disordered residues), we polish the obtained dataset (Figure 1A, Methods Figure 1). We extend an IDR to the N-, or C-terminus when there are less than 8 residues from the terminus of the protein to the nearest IDR. Further, when there are less than 8 residues in between two IDRs, we merge these two IDRs, including the residues in between, forming a single longer IDR. Next, if an IDR is separated by 8-16 residues from a terminus, or 8-16 residues from another IDR, and these residues adopt little secondary structure or have an average pLDDT below 70, we extend the IDR to the termini, or merge the two IDRs. The details of this data polishing procedure can be found in the methods section.

**Figure 1.**
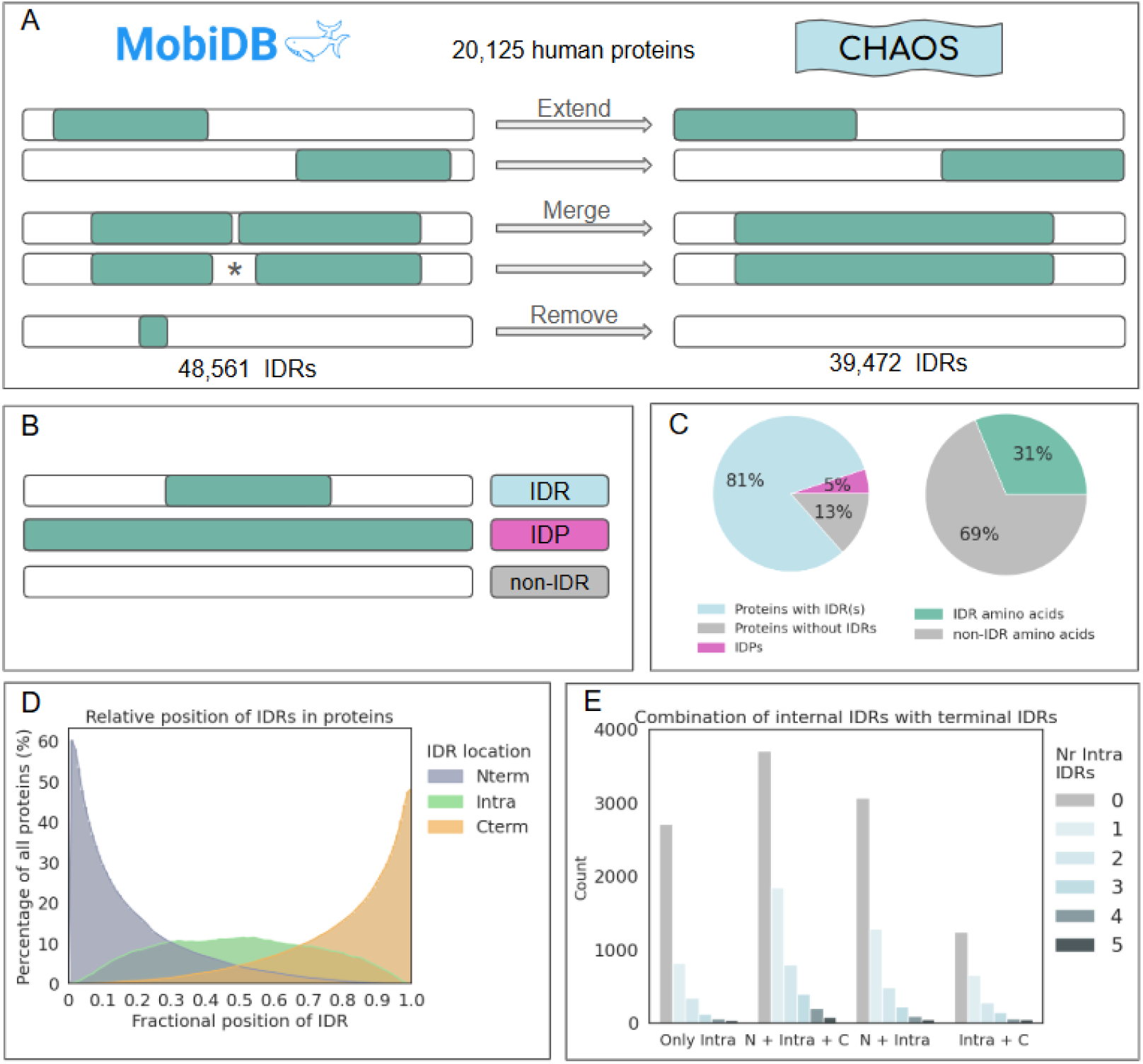
IDRs in human proteins. A) Schematic overview of how the IDRs obtained from MobiDB were extended, merged, or removed resulting in our CHAOS dataset, with 20,125 proteins and 39,472 IDRs; *:merged under special conditions. B) Schematic representation of the three protein categories considered in this work: IDR containing proteins, Intrinsically Disordered Proteins (IDPs), proteins without IDRs (non-IDR proteins). C) Left: Abundance of IDR containing proteins, proteins without IDRs and IDPs in this work. Right: The IDRs account for 31% of all amino acids in the dataset. D) Relative positioning of IDRs in all proteins. E) Prevalence of combinations of IDR location in proteins with and without IDRs. Only intra: protein with only internal IDRs, N + Intra + C: protein with an N-terminal, internal and C-terminal IDR, N + Intra: protein with an N-terminal and internal IDR, Intra + C: protein with an internal and C-terminal IDR. The colors indicate the number of internal IDRs.

### Prevalence and location of IDRs in human proteins

The resulting dataset contains 39,472 IDRs in 20,125 human proteins. From this, we define three protein categories (Figure 1B): IDR-containing proteins, with one or multiple IDRs and one or multiple non-IDRs are the majority (81%); Intrinsically Disordered Proteins (IDPs), that contain a single IDR that spans the entire protein, are 5%; and the proteins contain no IDRs at all are 13%. In total, the disordered residues account for 31% of all amino acids (Figure 1C).

More than half of all IDRs (57%) are located at protein termini: 32% at the N-terminus, and 25% at the C-terminus. The end-result of this distribution, is that most proteins (62%) contain an N-terminal IDR, about half (48%) contain a C-terminal IDR, and about three quarters (74%) of all proteins contain either N-or C-terminal IDRs. (Figure 1D). The random probability that an IDR with an average length (80 residues) is positioned in either the first or the last 8 residues of a protein with average length (524 residues) would be around 4% ((8/(524-80))*2). This is substantially lower than the mentioned 74%, suggesting that proteins are highly biased towards having IDRs at their termini. Noteworthy is that some terminal IDRs span nearly the entire protein (Supplemental Tables S1 and S2).

On the other hand, internal IDRs (42%) are less prevalent. Less than half (42%) of the proteins have one or more internal IDR (Figure 1D). The idea that internal IDRs could be viewed as linker regions3, suggests that proteins with internal IDRs are longer than proteins without internal IDRs. Indeed, proteins containing internal IDRs (average 741 residues) are considerably longer than proteins with N- or C-terminal IDRs (average 551 and 574 residues, respectively) (Supplemental Figure S1A). We find a correlation above the diagonal between the number of disordered residues and protein length (Pearson correlation coefficient 0.82) (Supplemental Figure S1B), suggesting long proteins contain relatively more disorder. This is mainly affected by more (Supplemental Figure S1C), rather than longer internal IDRs (Supplemental Figure S1D). However, with an average of two IDRs per protein and the bias to terminal IDRs in proteins, a protein is unlikely to contain multiple internal IDRs.

Indeed, the most frequently occurring combination of IDRs in a protein are an N- and a C-terminal IDR together, without internal IDRs (Figure 1E). The second most abundant group are proteins containing only an N-terminal IDR, while the third most common combination are proteins containing both N-, C-terminal and one internal IDR. Every internal IDR added to any of these categories reduces the number of proteins with that combination.

### Where in the human cell do we find disorder?

To study whether the amount of protein disorder varies at different subcellular locations, we incorporated UniProt-annotated subcellular locations and categorized each protein into different categories: plasma membrane, cytoplasm, secreted, cortex, nucleus, single organelle, multiple organelles or unknown localization (see methods for details) (Figure 2A). Note that each protein has been assigned to a single category, but in reality, proteins can shuttle between multiple cellular locations^37^.

**Figure 2.**
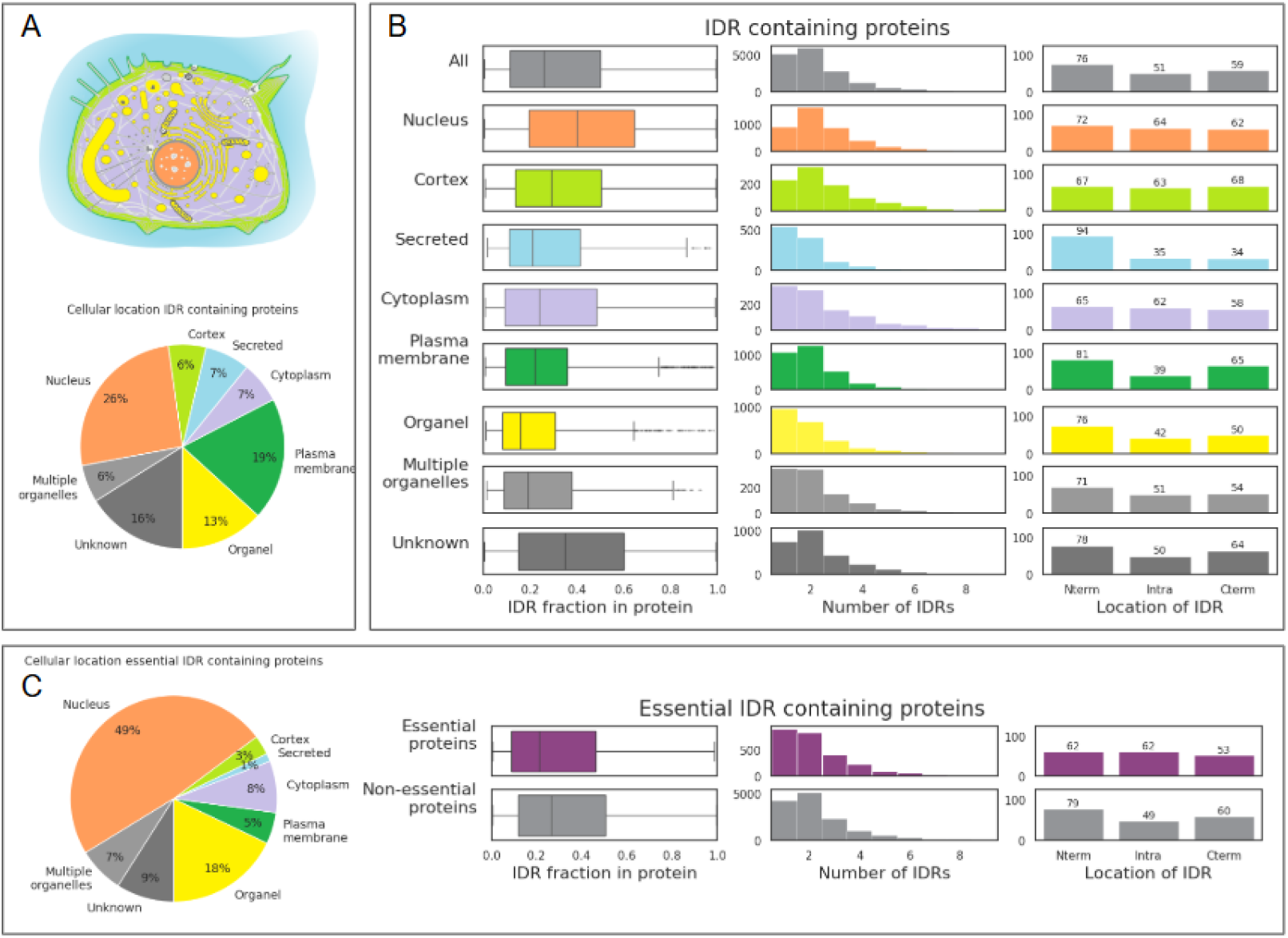
Cellular localization of IDR containing proteins. A) Graphic representation of the cellular locations defined for this work, wherein each color indicates a category (background figure obtained from https://www.swissbiopics.org/name/Animal_cell). The pie plot shows the distribution of proteins amongst these cellular locations. B) Boxplots of the fraction of IDR residues in IDR containing proteins (left). Distribution of the number of IDRs in IDR-containing proteins, wherein the y-axis represents the number of proteins (middle). The percentage (y-axis) of IDR containing proteins that contains an N-terminal (Nterm), internal (Intra) and C-terminal (Cterm) IDR (right). The row ‘All’ contains all IDR containing proteins, the following rows hold IDR containing proteins found at the mentioned cellular location. C) Cellular location of essential IDR containing proteins (left). Comparison of essential and non-essential IDR containing proteins for the fraction of IDR residues in the proteins, the number of IDRs in the protein and where the IDRs locate, similar as in B.

Proteins at different subcellular locations have different fractions of disorder (Figure 2B). Cortical and cytoplasmic proteins contain similar amounts of disorder (31-33%), while organellar, secreted and plasma membrane proteins contain less disorder (22-29%). Proteins in the nucleus (42%) and proteins without a known location contain more disorder (44%). The latter is likely due to experimental difficulties with highly disordered proteins. Within the organelle category we also find substantial differences in fractions of disorder (Supplemental Figure S2A). For example, endosomes contain a higher fraction of disorder (30%), while e.g. peroxisomes (20%), lysosomes (16%) and mitochondrial (21%) proteins have less disorder. It is possible that proteins in peroxisomes and lysosomes benefit from less disorder to prevent proteolysis, as an alternative to glycosylation. The low disorder in mitochondria might be related to their evolutionary descent from bacteria, which contain less protein disorder than human proteins^3^.

When investigating the number of IDRs per protein (Figure 2B), we observe that proteins located at the cortex, cytoplasm and nucleus contain more IDRs per protein (on average 2.8, 2.7, 2.6 IDRs per protein, respectively) than the average IDR containing protein (2.4 IDRs per protein). Secreted proteins contain the least IDRs per protein (1.9 IDRs per protein respectively), while proteins in organelles and at the plasma membrane are just slightly below average (2.1 and 2.2 IDRs per proteins, respectively). The protein length at different subcellular locations also differs, for example, cortical proteins are particularly long (Supplemental Figure S2B).

These data show that protein characteristics at different subcellular locations contribute to a unique environment at their specific locations. For example, long, highly disordered cortical proteins form a uniquely disordered environment just below the cell surface. The high amount of disorder at the cell cortex might be important for sensing and remodeling membranes^38–40^.

#### Cellular location determines IDR location

Most secreted and plasma membrane proteins contain N-terminal IDRs (94%, 81%, respectively) (Figure 2B). The high likelihood for N-terminal IDRs is likely due to the annotation of N-terminal localization peptides as IDRs. Such peptides are required to secrete proteins outside the cell or to translocate proteins into cell membranes. As localization peptides are often proteolytically removed^41^, these numbers likely do not reflect the functional composition of IDRs in secreted and plasma membrane proteins, but rather the complete protein before any processing. Further, C-terminal IDRs are less prevalent in secreted proteins (34%), while C-terminal IDRs are more prevalent in plasma membrane proteins (65%). Secreted and plasma membrane proteins both contain less internal IDRs (35%, 39%). Such proteins might render secreted and cell membrane proteins susceptible to proteolysis, altering their function or regulating activity levels.

Proteins that localize at the cytoplasm or cortex contain fewer N-terminal IDRs (65%, 67%) (Figure 2B). In turn, these proteins are enriched in either or both C-terminal or internal IDRs. Cytoplasmic and cortical proteins are enriched for internal IDRs (62%, 63%), while cortical proteins are enriched in C-terminal IDRs (68%). From these data it becomes clear that the location of the IDR in the protein is biased, which is reflected in differences per cellular location and protein length. Whether this is cause or causality remains unclear.

#### Essentiality

The essentialome is the collection of genes or proteins that are required for survival and proliferation in cells, highlighting fundamental pathways. Note that essentiality annotations we used stem from studies performed in single cell lines^29,30^ and might differ from essentiality in whole organisms. In our dataset we include 3,440 (17% of total) essential proteins, of which half (49%) are found in the nucleus (Figure 2C). Essential proteins that contain IDRs, have a similar fraction of disorder (30%) as non-essential IDR containing proteins (34%) (Figure 2C, Supplemental Figure S2C). Also, the average number of IDRs (2.4) does not substantially differ from non-essential IDR containing proteins (2.3) (Figure 2C, Supplemental Figure S2C). Note that essential proteins are approximately equally likely to contain a N-terminus, C-terminus or internal IDR (62%, 62%, 53%) (Figure 2C).

### What are intrinsic properties of IDRs?

#### How long are IDRs?

The length of IDRs might influence their function and properties^42–45^. On average, IDRs are 80 residues long (median 36 residues), which is shorter than (folded) regions between IDRs (average 189, median 118 residues) (Figure 3A). Overall, N-terminal (average 75, median 35 residues) and internal IDRs (average 75, median 34 residues) are of similar length, while C-terminal IDRs are longer (average 97, median 43 residues) (Supplemental Figure S3A). At different cellular locations we observe that proteins located at the cortex or in the nucleus contain the longest IDRs, while secreted, organellar (not nuclear) and membrane proteins contain the shortest IDRs. Furthermore, nuclear proteins have longer, and secreted proteins have shorter N-terminal IDRs than other locations. Cortical C-terminal IDRs are the longest while the length of internal IDRs is relatively constant at different locations (Supplemental Figure S3B).

**Figure 3.**
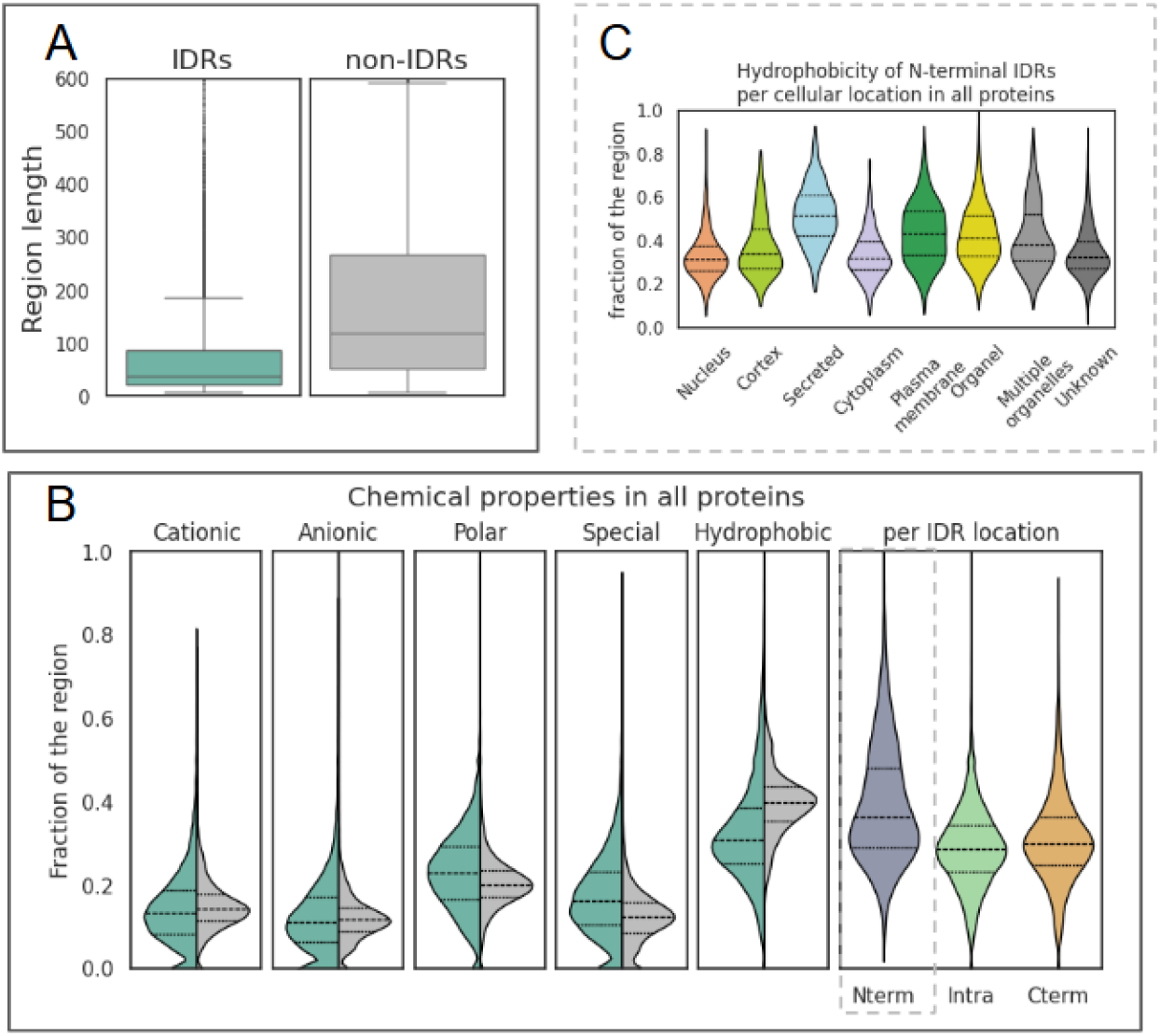
Properties of IDRs and non-IDRs. A) Length in the number of residues for IDRs and non-IDRs in our dataset. B) Chemical properties of IDRs and non-IDRs divided into cationic (R,H,K), anionic (D,E), polar (S,T,N,Q), hydrophobic (A,V,I,L,M,F,Y,W) and special (C,P,G) residues. Hydrophobicity is also shown per IDR location in the protein (Nterm: N-terminal, Intra: internal, Cterm: C-terminal IDR). C) Hydrophobicity of N-terminal IDRs found in IDR containing proteins at different cellular locations.

#### IDRs are not hydrophobic

The chemical composition of IDRs and non-IDRs shows a different profile (Figure 3B). IDRs are less hydrophobic and slightly enriched in polar and special residues compared to non-IDRs, while cationic and anionic (charged) residues occur in approximately equal fractions. These findings are in line with other studies^46–48^. When investigating the chemical properties of IDRs based on their position in the protein, we find that N-terminal IDRs have the most distinctive chemical profile (Supplemental Figure S4). These IDRs are considerably more hydrophobic than internal and C-terminal IDRs. The increased hydrophobicity could be due to localization peptides, which are known to contain a hydrophobic region. Such localization peptides are used to direct proteins to organelles, into the secretory pathway and translocate membrane proteins into the membrane^49^. Accordingly, we find increased hydrophobicity of N-terminal IDRs in secreted, organellar and plasma membrane proteins (Figure 3C).

#### Effects of missense variants

Single amino acid changes can affect protein folding activity and stability, causing pathogenic phenotypes. Pathogenic missense mutations in the context of ordered regions in proteins are well-studied and often well-understood. To understand if missense mutations in disordered residues are similarly pathogenic as mutations in ordered residues, we incorporated data from AlphaMissense^50^ into the CHAOS dataset. AlphaMissense assigns a pathogenicity score (0-1) for every possible mutation, which are classified as likely benign, likely pathogenic, or uncertain.

The distribution of the pathogenicity scores in IDRs and non-IDRs is profoundly different (Figure 4A). Most missense mutations in IDR residues (62%) are likely benign, but one in five (19%) are likely pathogenic. Although this percentage may seem small, it should not be overlooked, as it implies that these residues can play crucial functional roles. In contrast, missense mutations in non-IDR residues are mostly likely pathogenic (54%), and less often likely benign (26%). Missense mutations in aromatic residues (F, Y, W) are more likely pathogenic than other amino acids, both in IDRs and non-IDRs (Figure 4B, Supplemental Figure S5). The median pathogenicity in IDRs of these amino acids together with lysine, methionine and isoleucine is uncertain. All other missense mutations in IDR residues are likely benign. Noteworthy are missense mutations in cysteine residues, that are almost exclusively likely pathogenic in non-IDRs but likely benign in IDRs. Missense mutations in histidine, threonine and proline IDR residues are almost exclusively likely benign. Lastly, missense mutations in glutamines are almost all likely benign, independent whether these are IDR or non-IDR residues.

**Figure 4.**
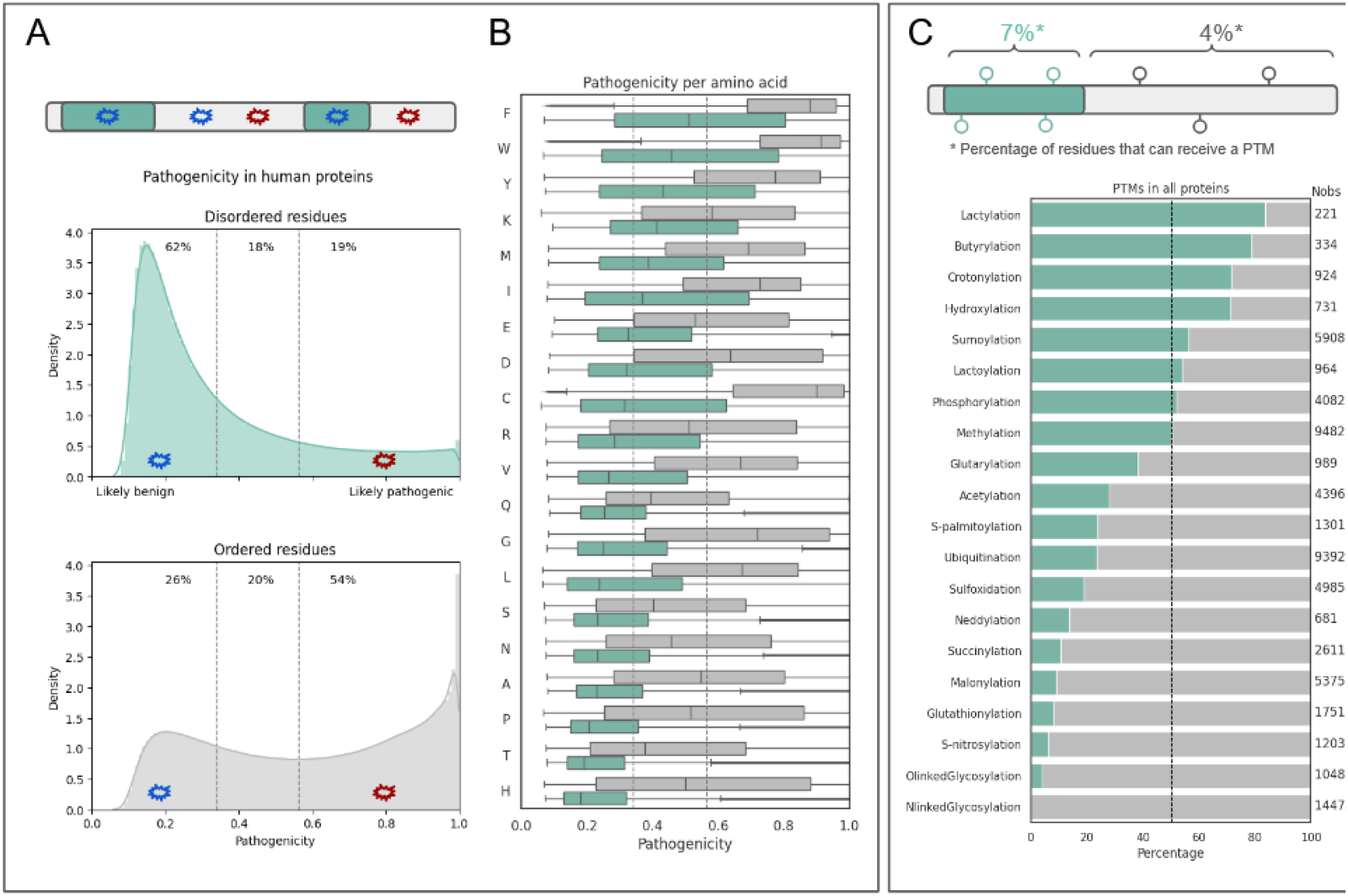
A) Cartoon representation of likely pathogenic (red) and likely benign (blue) missense mutations in IDR and non-IDR regions (top). Histograms representing the pathogenicity of disordered residues (green) and ordered residues (grey). The disordered and ordered residues contribute to 31% and 69% of all residues, respectively. The area under both curves is set to 1. Cut-offs for likely benign (left, blue) and likely pathogenic (right, red) are indicated with a grey dotted line; the limits are set according to AlphaMissense50. B) Pathogenicity per amino acid in IDR residues (green) and non-IDR residues (grey) represented as boxplots. C) Schematic representation of PTMs sites in a protein, wherein a residue can obtain a single or multiple PTMs (top). Relative presence of PTMs in IDRs vs. non-IDRs for all proteins in the dataset (bottom). The dotted vertical lines mark the 50% border, and the numbers on the right-hand sides represent the number of PTMs found in total for that bar. For clarity only PTM types that were found >=200 times in our dataset are shown.

#### Disordered residues are enriched in PTMs

Post-translational modifications (PTMs) can create unique chemical profiles in proteins, while exerting important regulatory functions^51,52^. To study PTMs in human proteins, we first mined PTM sites from MobiDB. MobiDB contains a small set of well-annotated modifications based on UniProtKB^31^, and additional phosphorylation sites from the Scop3P database^53^. We enriched this set including PTMs from dbPTM^32^, a database containing nearly 80 PTM types integrated from over 40 databases. dbPTM contains manually curated and experimentally validated PTM sites, as well as PTMs curated by literature and predicted PTM sites. Combining all PTMs found in human proteins in the CHAOS dataset, resulted in 610,446 PTMs at 512,671 unique amino acids, covering more than 60 different PTM types.

Similar to other studies^54^, we find that residues that can be modified are relatively more prevalent in disordered residues (7%) than in ordered residues (4%) (Figure 4C). This could be explained by surface accessibility, amino acid proportion, or the regulatory role of IDRs. The average number of PTMs in our dataset at a particular residue is similar in disordered residues and ordered residues (1.1 and 1.3 PTMs per PTM residue, respectively). Be aware that more PTMs and/or PTM residues in either ordered or disordered residues might be found in the future.

The most enriched modifications in disordered residues relative to ordered residues are lactylation (polar), butyrylation and crotonylation (hydrophobic modifications) (Figure 4C). All these modifications occur on lysines and fall under the acylation umbrella^55^. The fourth most enriched PTM type in IDRs is hydroxylation. The latter seems particularly enriched in secreted proteins (Supplemental Figure S6), where it might aid the formation of supramolecular structures and lower the susceptibility to proteolysis^56,57^. Notably, N-linked glycosylation has not been detected in IDR residues. Essential proteins contain relatively more IDR PTM sites (11%), which is in line with a previous study that shows more PTMs are found in essential proteins^58^.

The most prevalent PTM in our dataset is phosphorylation, which is not clearly enriched in disordered nor ordered residues (Figure 4C). However, phosphorylation on serine residues is overrepresented in IDRs, while phosphorylation on tyrosine residues is enriched in non-IDRs relative to IDRs. Threonine phosphorylation is not enriched in either category (Supplemental Figure S7). These results are in line with the relative occurrence of these three residues in either IDRs or non-IDRs (Supplemental Table S3). Of note, both acetylation and ubiquitination are considerably enriched in non-IDRs compared to IDRs. Since ubiquitination can lead to degradation, the latter could indicate IDRs follow different degradation pathways than non-IDRs. This difference possibly is an important factor in cell homeostasis^59^.

### What have we learned from protein structure prediction methods?

To capture the discrepancies between the predicted protein structure models in the AlphaFold database^22,36^ and our data, we calculated the percentage of residues that adopt a secondary structure, and the average pLDDT value (reflecting the confidence of the model) for each IDR and non-IDR (Figure 5A). As expected, we find lower pLDDT values and lower percentages of residues that adopt a secondary structure for IDRs than for non-IDRs (Figure 5B). Typically, the pLDDT of IDRs is low (39 on average), as expected. However, rather surprisingly, 43% of residues in IDRs adopt a secondary structure in predicted models. This is considerably higher than the naive expectation that IDRs contain 0% secondary structure. Strikingly, AlphaFold predicts some IDRs to be completely ordered. When we select such regions, we find 474 proteins containing 587 IDRs with a high percentage (>92%) of secondary structure in total, of which 84% is an internal IDR. Based on cellular localization, IDR and non-IDR length, and chemical properties, they are not similar to non-IDRs (Supplemental Figure S8).

**Figure 5.**
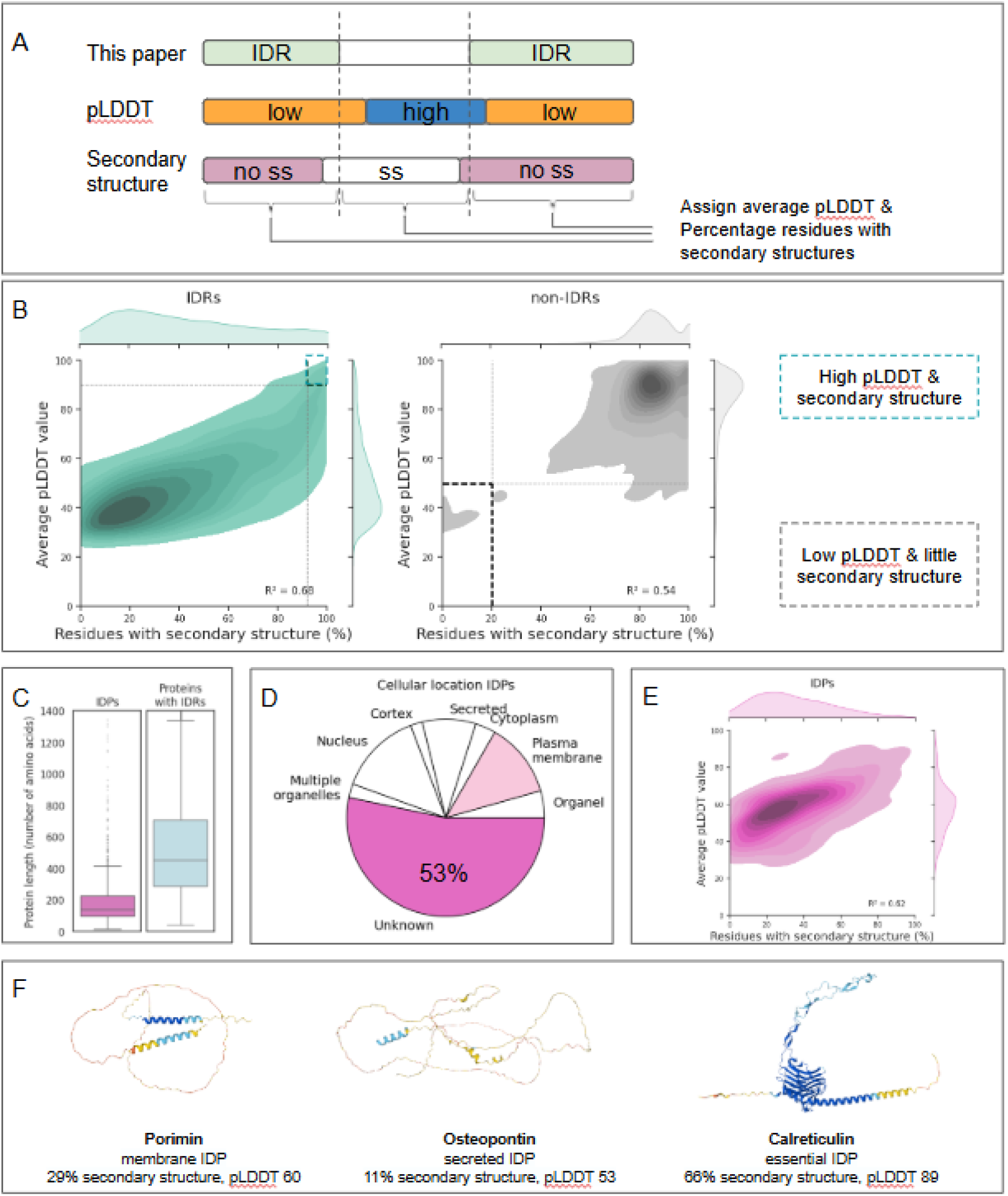
IDRs and IDPs are not fully disordered. A) Schematic representation of IDRs in a protein sequence. The pLDDT score of the corresponding AlphaFold model and the percentage of residues with secondary structure do not perfectly align to the CHAOS dataset. Average values are annotated per region. B) Correlation between the percentage residues that adopt secondary structure and the average pLDDT score of the IDRs (left) and non-IDRs (right) found in our dataset. R2 marks the Pearson correlation coefficient. The dotted lines in the IDR plot (pLDDT >=90, percentage of residues that adopt secondary structure >=92%) mark IDRs with high confidence and high percentage (above the third quartile) of secondary structure. The dotted lines in the non-IDR plot (pLDDT <=50, percentage of residues that adopt secondary structure <20%) mark non-IDRs with low confidence and little (below the first quartile) secondary structure. C) The length of IDPs and IDR containing proteins. Y-axis represents the length in number of amino acids. D) The cellular localization of IDPs in our dataset. E) Correlation between the percentage residues that adopt secondary structure and the average pLDDT score of IDPs. R2 marks the Pearson correlation coefficient. F) 3D representation of example cases of IDPs to illustrate IDPs are not predicted to be fully disordered. Coordinates originate from the AlphaFold database (Porimin: ID Q8N131, Osteopontin: ID P10451, Calreticulin: ID P27797), and the models are colored by pLDDT (dark blue: pLDDT >90, light blue: 90 > pLDDT > 70, yellow: 70 >= pLDDT >50, orange pLDDT <=50).

Next, we investigated the pLDDT and percentage of residues that are predicted to adopt a secondary structure in non-IDRs. Surprisingly, AlphaFold predicts some non-IDRs with low pLDDT and a low percentage of residues that adopt secondary structure. Selecting these seemingly disordered non-IDRs, we find 449 non-IDRs in 402 proteins, which are enriched in N-terminal regions (38%). When we compare the chemical properties the non-IDRs with low pLDDT and a low percentage of residues that adopt secondary structure, we observe similarities to the chemical properties of IDRs. Therefore these non-IDRs are somewhat like IDRs and could therefore be incorrectly annotated (Supplemental Figure S8).

### Are IDPs more ordered than they should be?

In this work, we define Intrinsically Disordered Proteins (IDPs) as proteins that contain a single IDR spanning from the first to the last residue (after merging and expanding MobiDB IDRs). This strict annotation results in 1017 (5%) IDPs. These proteins are on average 201 residues long (Figure 5C), which is considerably longer than the average IDRs (80 residues), but considerably shorter than average proteins (524 residues).

More than half of the IDPs (53%) lack subcellular location annotation (unknown) (Figure 5D). This is likely due to experimental difficulties and underlines the importance of a broad characterization. Strikingly, we also find a considerable number of IDPs located in the plasma membrane (12%). Membrane proteins typically contain at least one transmembrane spanning helix (typically 20 amino acids long^60^), which implies IDPs in the plasma membrane may not be fully disordered at all times. Most likely such proteins form β-barrels, or similar secondary structures that only exist when multiple strands are lined up. This implies that membrane-IDPs should be considered in their native environment and with their native stoichiometry. It is also possible that protein disorder may enhance the fluidity of the plasma membrane.

We propose defining IDPs as a distinct class of proteins, with membrane IDPs as a specific subclass. To our knowledge only MINAR-like proteins (e.g. P59773 and Q9UPX6) have been characterized as a distinct membrane-IDP protein family in Pfam, however IDPs and membrane IDPs as such, have not been classified^61^.

Although the chemical properties of IDPs and IDRs are more similar to each other than to those of non-IDRs, IDPs contain relatively more special residues (C,P,G) (Supplemental Figure S9A). We also find less PTM sites on IDPs (5%) than in IDRs (7%). This may be due to a lack of studies on IDPs, like the absence of cellular localization annotations. Further, we note that only 1% (44/3440) of essential proteins is an IDP, whereas 6% (973/16,685) of non-essential proteins is an IDP. This substantial difference suggests that combining the characteristics of essentiality and IDPs is evolutionary disadvantageous.

While IDPs are expected to be fully disordered, AlphaFold predicted that 35% of IDP residues adopt a secondary structure with an average pLDDT score of 58 (Figure 5E). Both values are higher for IDPs than for IDRs. The pLDDT and percentage of residues that adopt a secondary structure are even higher for plasma membrane IDPs (Supplemental Figure S9B). For example, Porimin, a membrane-IDP, contains helical parts accounting for 29% residues with secondary structure (Figure 5F). On the other hand, Osteopontin, a secreted IDP, looks more like the expected IDPs, having almost no secondary structure elements (11%) (Figure 5F). Exemplifying the complexity of IDPs, we highlight Calreticulin, an essential IDP that localizes at the endoplasmatic reticulum (ER), binding calcium ions. The protein is predicted with almost solely secondary structure elements (81%) (Figure 5F), although it is experimentally shown that the P-domain is unfolded in solution^62^. However, it is argued that Calreticulin might adopt different conformations in response to its environment^63^. These examples illustrate that while IDPs can be disordered on their own, they could adopt secondary structure elements depending on their biological environment. Thus, IDPs are a unique class of short proteins that display more secondary structure than their name suggests.

## Conclusion and Outlook

The knowledge of protein structures and structure-related functions is rapidly expanding. Despite the increasing attention for IDRs in proteins^64^, systemic characterization is lacking. Our analysis provides a unique view, as we combine publicly available data and predicted structures of human proteins to form the CHAOS dataset we present here, analyzing many aspects of ordered and disordered regions of human proteins.

Overall, we confirm that most human proteins contain one or more IDRs. Strikingly, terminal IDRs are a lot more prevalent than internal IDRs. We also find that proteins containing internal IDRs are considerably longer. Furthermore, the cellular location of a protein determines the preferred IDR location in the protein sequence. N-terminal localization peptides are likely annotated as IDRs. Therefore, it might be worth considering localization peptides as a subclass of IDRs. However, more detailed studies on differences and similarities of IDRs and localization peptides are complicated due to their wide variety^65^ and lack of experimental validation^66^. Intriguingly, IDRs themselves are enriched in certain PTMs. Somewhat expectedly, missense mutations in disordered residues are less likely to be pathogenic, than similar mutations in ordered residues, as it is likely that the latter would affect e.g. enzymatic function.

To characterize IDRs in more detail, our dataset could be further enriched with conservation scores. However, it is difficult to obtain consistent conservation scores for all human proteins. Namely, some proteins can be traced back to the last universal or the last eukaryotic common ancestor, while others are restricted to mammals, which complicates analysis. In addition, traditional alignment algorithms alone might not be particularly informative on IDRs^67^. Adding other protein characteristics like linear motifs, genetic diversity or conformational ensembles^68^, could aid to assign informative conservation scores to IDRs and their characteristics. Together these could lead to a more detailed classification of IDRs in proteins, which ultimately sheds light on how differences in IDR composition relate to functionality.

As IDRs have been proposed as the driving force of phase separation^44^, we identified phase separating proteins in our dataset based on annotations in MobiDB (which are based on the PhasePro^69^ and PhaSePro^70^ databases) with the intention to understand how their IDRs are different from IDRs that do not induce phase separation. We found that phase separating proteins contain a higher percentage of disorder and their IDRs are longer than proteins that are not annotated to phase separate (Supplemental Figure S10A and B). We also found relative enrichment of methylation in IDRs in phase separating proteins (Supplemental Figure S10C). Methylation reduces charge and increases the hydrophobicity and might be acting as an on/off switch to phase separating behavior^71^. However, our data contains only 289 phase separating proteins, which we do not consider sufficient to draw robust conclusions. Further we find 28 proteins without IDRs that are annotated to be involved in phase separation (Supplemental Table S4), which might be a glitch in our data. However, the latter proteins might have been annotated because the discrepancy between a protein driving phase separation or being in the phase separation bubble is not always clear.

Disordered regions are known to be able to adopt secondary structure depending on their biochemical context, e.g. proximity of another protein upon complex formation^72^. For example, IDPs can have a packing density and core hydration^68^, indicative of gaining order. There are also examples of IDPs adopting 3D structures^72^. Ensemble modelling indicates that IDRs have conformational properties that relate to protein function and cellular localization^34^. While only few conditionally folding IDRs are annotated, we find that almost half of all IDR residues adopt some secondary structure in predicted models. This is in line with a finding that AlphaFold has the tendency to predict the folded state of experimentally verified conditionally folding IDRs^26^. Another possibility is that AlphaFold has mistakenly predicted some structural elements (hallucination). As prediction algorithms have been trained on the folded state of proteins they tend to overestimate the structural propensity.

Collectively, our observations emphasize the complex nature of IDRs and IDPs in human proteins. This study, and many others, are considering IDRs or IDPs in single proteins. However, most proteins in the cell interact with proteins or other macromolecules. Current computational developments are capable of predicting the structure of higher order protein complexes^73^. This opens the possibility to systematically study e.g. sequence properties of IDRs, the role of PTMs in interface formation, or the nature of disorder-to-order transitions. Large language models and generative AI approaches might also prove useful to provide better annotations to IDRs an their properties in cells and relate them to physiological and disease states.

## Methods

### Data collection

The data of the human genome was downloaded from MobiDB^28^ on 2025-01-31 in JSON format. We chose MobiDB because it is user-friendly both for humans and computers and was recently updated (2022-07) at the time the project started. Swiss-prot annotated human protein information was downloaded from UniProtKB^31^ on the same day.

Dictionary of Secondary Structure in Proteins (DSSP)^35^ was used to define the secondary structure elements that AlphaFold2^22,36^ predicted for the human proteins (AlphaFold_human, Reference Proteome: UP000005640). The secondary structure elements were stored per residue. Other (“ ”) residues were defined as disordered residues, whereas Alpha-helix (H), Beta-bridge (B), Strand (E), Helix_3 (G), Helix_5 (I), Helix_PPI (P), Turn (T) and Bend (S) were considered as ordered residues. UniProt primary accession codes for which no AlphaFold prediction was available in the AlphaFold protein structure database (75 proteins) were deleted from the dataset, as no DSSP information could be retrieved. Additionally, the largest human proteins in the AlphaFold database (for which the coordinates are stored in multiple coordinate files because they are too large to fit into a single file, 208 proteins), were removed from the dataset as these are hard to handle technically. The UniProt primary accession codes of the removed proteins can be found in Supplemental Methods S1. This led to a final selection of 20,125 human UniProt primary accession codes.

For each of these proteins, we extracted the AlphaMissense data from the AlphaFold database using its API. For each protein, the average pathogenicity score was calculated for all possible mutations at every amino acid position. Subsequently, pairwise protein alignment was performed to reconcile the residue numbering between the AlphaMissense data and the MobiDB data, ensuring consistency across the datasets.

To be able to incorporate information about PTMs (post-translational modifications), the PTM annotations from MobiDB were stored and enriched with PTMs present in dbPTM^32^, downloaded on 2023-02-01]. From dbPTM we selected all PTMs that are found in human proteins that are in the 20,125 UniProt primary accession codes mentioned above. This resulted in a list of 610,446 PTMs.

### Dataset construction

To define IDRs 20,125 human proteins, the IDRs annotated in MobiDB were mined and stored with their start and end position for each protein. To avoid two IDRs with no or small sequence distance, the IDRs annotated by MobiDB were evaluated based on their length, the number of amino acids in between two IDRs, and the DSSP annotation of the secondary structure that AlphaFold2 predicts for the IDR regions. The predicted local distance difference test (pLDDT) score was used as a justification metric. Namely for an ordered region in between two IDRs with high pLDDT (>70%), it is most likely an ordered region^26,27^. On the other hand, if pLDDT of these residues is low (<70%), it is most likely a disordered region^26,27^, so we can merge the two IDRs with the amino acid in between them into a single IDR. Based on the average length of unmodeled regions in PDB-REDO structure models, we decided to keep a minimal IDR length of 8 amino acids. We also set this requirement, because 8 amino acids could form a two-turn helix, which we consider the smallest possible secondary structure element.

The decision-making process is as follows:

1. If there are equal to or less than 8 amino acids present between two IDRs, the two IDRs are merged with the amino acids in between into one single IDR when both IDRs are independently longer or equally long as the number of amino acids in between them.
2. If there are more than 8 up to including 16 amino acids between two IDRs, those IDRs will be merged with the amino acids in between into one single IDR when there are less than 80% ordered residues annotated by DSSP for the region in between and both IDRs are independently longer or equally long as the number of amino acids in between them. The two IDRs will also be merged when AlphaFold predicted more than 80% ordered residues being present in the region between the two IDRs, and this 80% holds at least 8 amino acids, and the average pLDDT value of the region between the two IDRs is below 70%. Again, merging only happens when the region in between and both IDRs are independently longer or equally long as the number of amino acids in between them.
3. The same logic as described in 1 and 2 is applied for the N-terminus to the first IDR in the protein, and between the end of the last IDR in a protein to the C-terminus.
4. If an IDR is shorter than 8 amino acids in length, it is removed from the dataset.

If the above criteria are not met, IDRs will not be merged nor deleted and kept as annotated by MobiDB. This process is schematically described in Methods Figure 1.

**Methods Figure 1:**
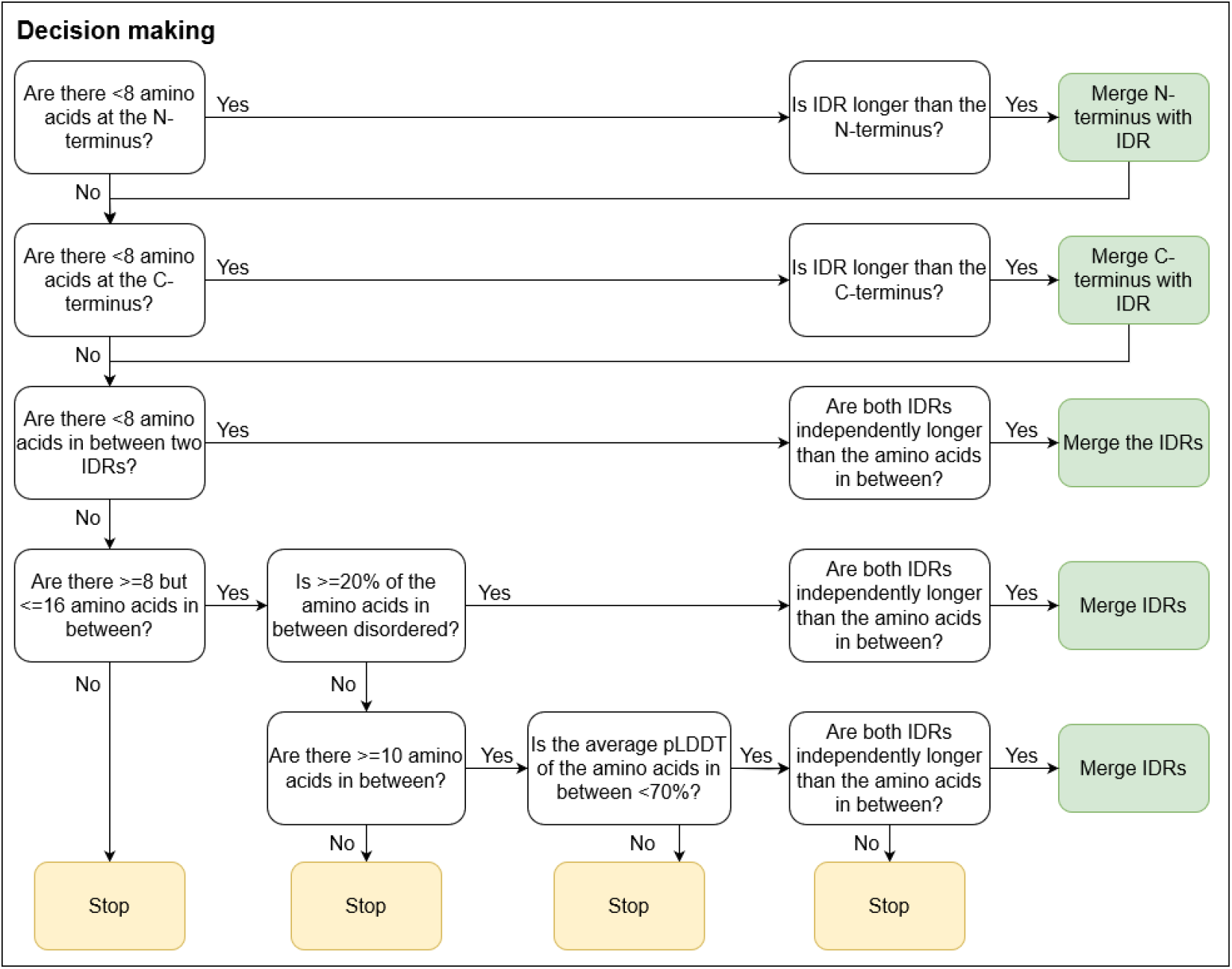
Decision tree for extending and merging IDRs.

### IDR dataset

For each of the unique UniProt primary accession codes in the dataset, several parameters and characteristics of the human proteins were mined or calculated; a summary is in Methods Figure 2.

**Methods Figure 2:**
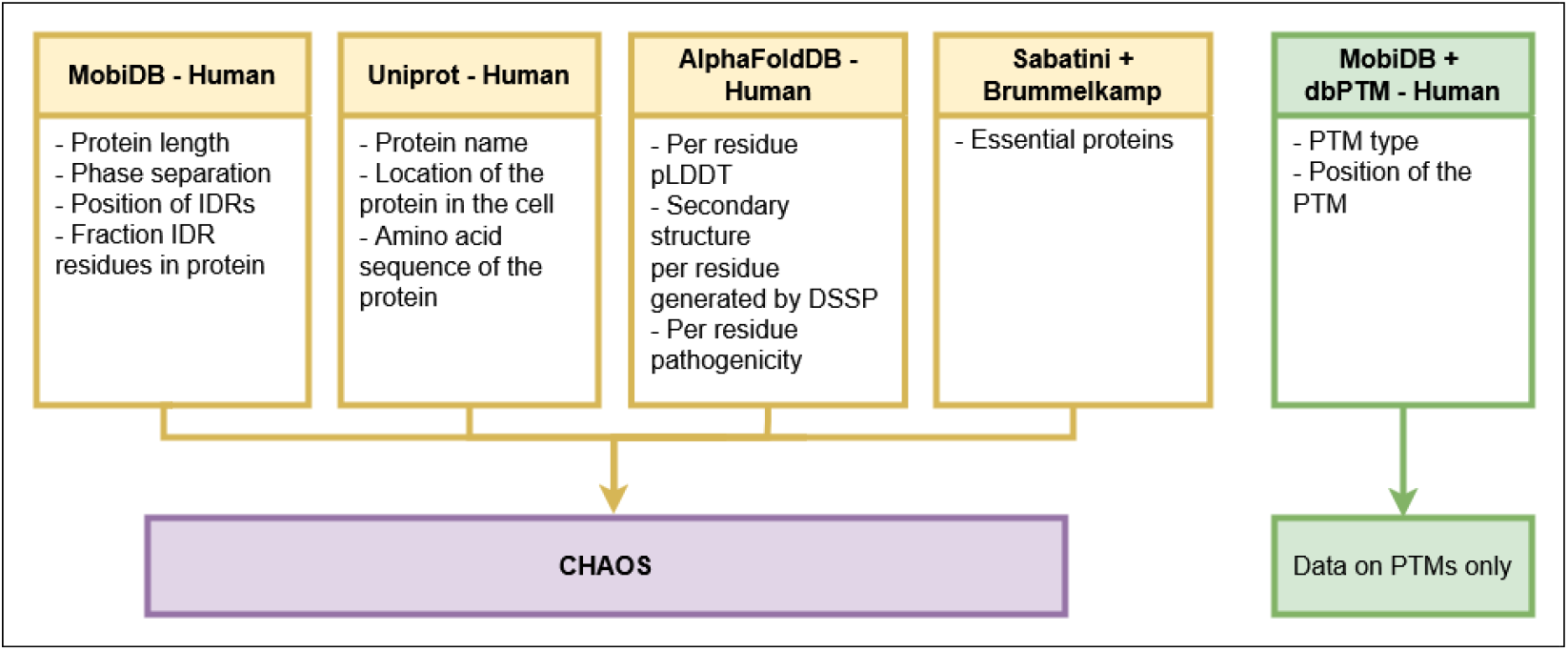
Summary of the data obtained for human proteins, their IDRs and characteristics.

#### Protein characteristics - length, IDR presence and cellular localization

On the protein level, for each UniProt primary accession code we state the protein length, the name of the protein as annotated in UniProt, the number of IDRs in the protein and the number of IDR residues. We calculated the IDR fraction per protein by dividing the number of IDR residues in a protein by the protein length. Additionally, we defined the cellular localization of each protein based on the annotation found in UniProt. We define the categories:

- Nucleus: proteins that localize at the nucleus.
- Organelles: proteins that localize to mitochondria, endoplasmic reticulum, endosome, golgi, autophagosome, phagosome, peroxisome, melanosome, lysosome or microsome.
- Multiple organelles: Proteins that localize to more than one of the organelles.
- Cortex: proteins that localize to places in the cell that are involved in sensing the environment, but do not localize to organelles. This category entails locations such as gap junctions, cilia, protrusions the cell cortex, collectively called interactions.
- Plasma membrane: transmembrane proteins and membrane-anchored proteins. Note that this category does not include proteins in the membranes of organelles, those are classified into the organelles.
- Secreted: Proteins that are extracellular.
- Cytoplasm: Proteins that are annotated to localize in the cytoplasm *and do not fall* in any of the above categories.
- Unknown: proteins without any known cellular location annotation.
- Miscellaneous: proteins that have a cellular location, but do not belong to any of the above categories (9 proteins). These proteins are not shown in further analyses.

We mark the proteins involved in phase separation based on the in MobiDB integrated annotations for this feature, which are originally retrieved from PhaSepDB^69^ and PhaSePro^70^. In order to determine if a protein is associated with an essential gene, the genes marked as essential in one or more cell-lines by Blomen et al.^29^ or Wang et al.^30^ were mapped to corresponding UniProt identifiers using the UniProt IDmapping tool [https://www.uniprot.org/id-mapping].

Pairwise Alignment of the protein sequences of the different source databases (Uniprot, AlphaFoldDB, dbPTM) were mapped to our main source database (MobiDB) to ensure consistency while comparing amino acid positions and types. To score the number of mismatches and gaps in protein alignments, the BLOSUM62 substitution matrix was used in combination with default scoring settings as in BlastP^74^.

#### IDR and non-IDR characteristics

For each IDR and non-IDR region we annotated the location of the region in the protein based on their amino acid positions. Regions starting at the first residue of the protein are annotated as N-terminal (Nterm) regions, regions that end at the last residue of the protein are marked as C-terminal (Cterm) regions. Regions that start and end at any location that is not the first or last residue of the protein, are marked as internal (Intra) regions. Proteins that contain no IDR residues are marked as ordered protein (OP) and proteins solely containing IDR residues are annotated as IDPs.

For each IDR and non-IDR region of the proteins in our dataset, we calculated the average pLDDT score based on the predicted protein structures by AlphaFold2. Additionally, we define the fraction of ordered residues in the region. For this purpose, we characterized the amino acids into five main categories: cationic (R,H,K), anionic (D,E), polar (S,T,N,Q), hydrophobic (A,V,I,L,M,F,Y,W) and special (C,P,G) (Supplemental Table S3). For each region, we counted how many residues occur in each of the categories and calculated their relative percentage by dividing this count by the total of amino acids in the region.

### Final dataset

The procedures above resulted in a final IDR dataset containing 20,125 proteins with 39,472 IDRs in total. In this set 1017 (5%) UniProt primary accession codes are annotated to be IDP (100% disordered), 16,396 (81%) have at least one IDR, and 2712 (13%) proteins have no IDRs annotated.

### Analyses

The datasets were loaded using Pandas 2.0.1 [https://zenodo.org/records/10957263] and plots were generated using Seaborn 0.13.2, all using Python 3.11.3. Box-, violin- and kde- (correlation) plots were created with a minimum of 30 data points^75^, if the desired sub-dataset was smaller, a stripplot was made to represent the data more realistically. Mean and median values are calculated for datasets containing at least 5 observations^75^.

### Used software and packages

BioPython 1.81 ^76^ ; DSSP 4.3 ^35^ ; Numpy 1.24.3 ^77^ ; Pandas 2.0.1 ^78^ ; Pdbecif 1.5 ^79^ ; Python 3.11.3 ; Seaborn 0.13.2 ^80^

## Contributions

Conceptualization: JB IdV DAS RB RX MK AP Methodology: IdV DAS JB RB RX MK Formal analysis: JB IdV DAS Investigation: JB IdV DAS Data Curation: IdV DAS Writing - Original Draft: JB & IdV (all), DAS (methods) Writing - Review & Editing: DAS, RB, BE, MU, MK, RX, AP Visualization: JB & IdV Resources and Funding acquisition: AP

## Acknowledgements

The scope and direction of this research were collectively shaped and managed by all PhD candidates from the lab of the last author (AP) and carried out collaboratively, part-time, without the usual formal supervision. This fostered scientific independence and creativity, resulting in a dynamic, self-driven research process. The project strengthened teamwork and problem-solving skills and demonstrated the potential of such research structures.

We would like to express our gratitude to our department’s freedom, scientific interest and thoughtful comments, that all contributed to the shaping of this project. Specifically, we have had fruitful discussions on diverse implications of this study with Robbie P. Joosten, Hans Wienk, Maarten L. Hekkelman, Shun Lee, Felipe R. Rodriguez, Gian-Luca McLelland and Tim Schröder. Further, we like to thank Boğaç Erçığ, Michael Uckelmann and Titia Sixma for reviewing the text. We thank the Research High Performance Computing facility of the Netherlands Cancer Institute for providing and maintaining computation resources.

