## Supplemental for "Disentangling the CHAOS of intrinsic disorder in human proteins"

### Supplemental information

Supplemental Figure S1: A) Protein length of proteins containing no IDRs, proteins containing IDRs, and of proteins containing an N-terminal (Nterm), at least one internal (Intra) or a C-terminal (Cterm) IDR. B) The protein length and the number of IDR amino acids in the protein correlate; Pearson correlation coefficient: 0.70. C) Proteins with increasing number of internal IDRs are longer. For clarity proteins with more than 11 internal IDRs are not shown. D) The length of individual IDRs does not correlate with the protein length; Pearson correlation coefficient: 0.20.

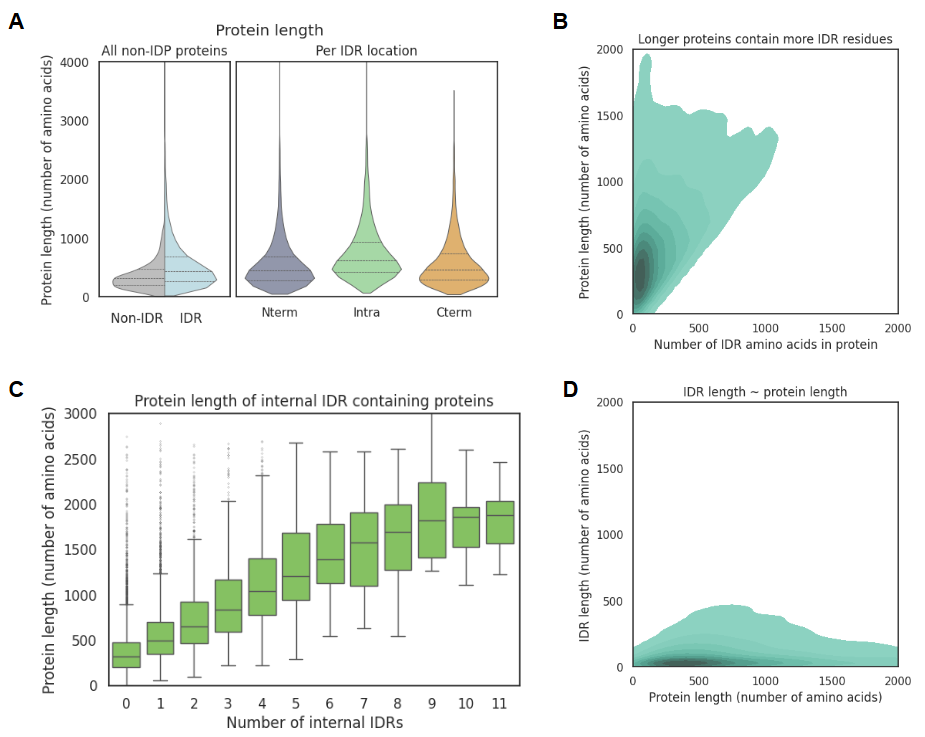

Supplemental Figure S2: A) Number of found proteins, protein length, IDR fraction and number of IDRs for IDR containing proteins found in organelles. B) Protein length of IDR containing proteins for all (grey) and per cellular localization. C) Fraction of IDR residues and the number of IDRs in essential IDR containing proteins for all (grey) and per cellular location.

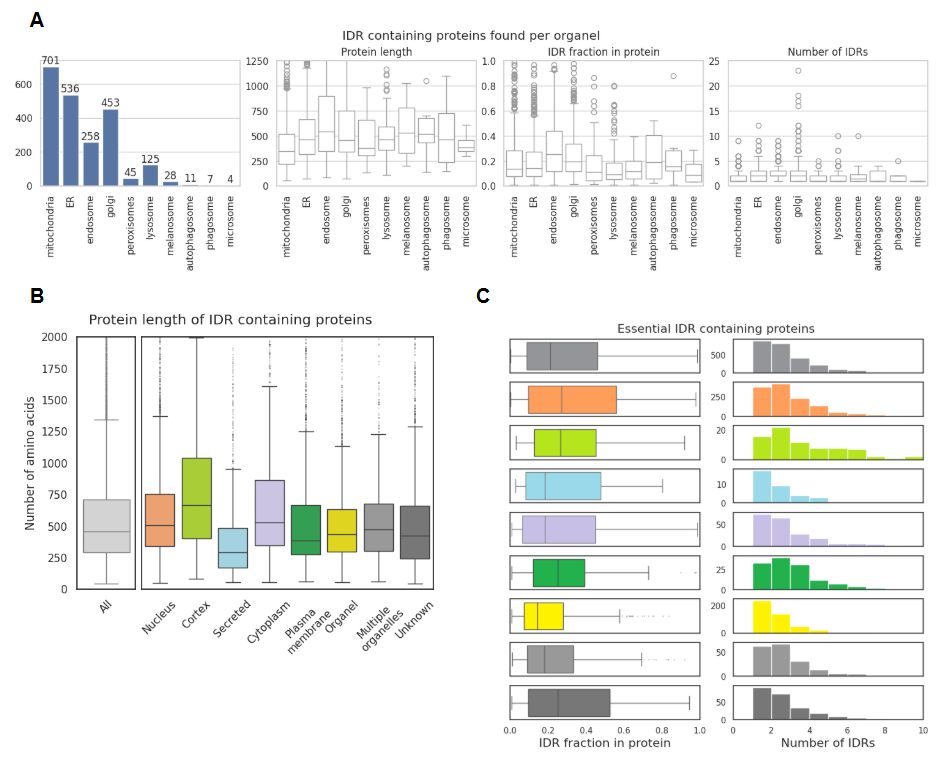

Supplemental Figure S3: A) Length of IDRs (left, green) and length of IDRs located at the N-terminus (Ntrem, purple), internally (Intra, light green), or C-terminus (Cterm, orange). B) Length of IDRs (left, white) and length of IDRs located at the N-terminus (Ntrem, purple), internally (Intra, light green), or C-terminus (Cterm, orange) per cellular location.

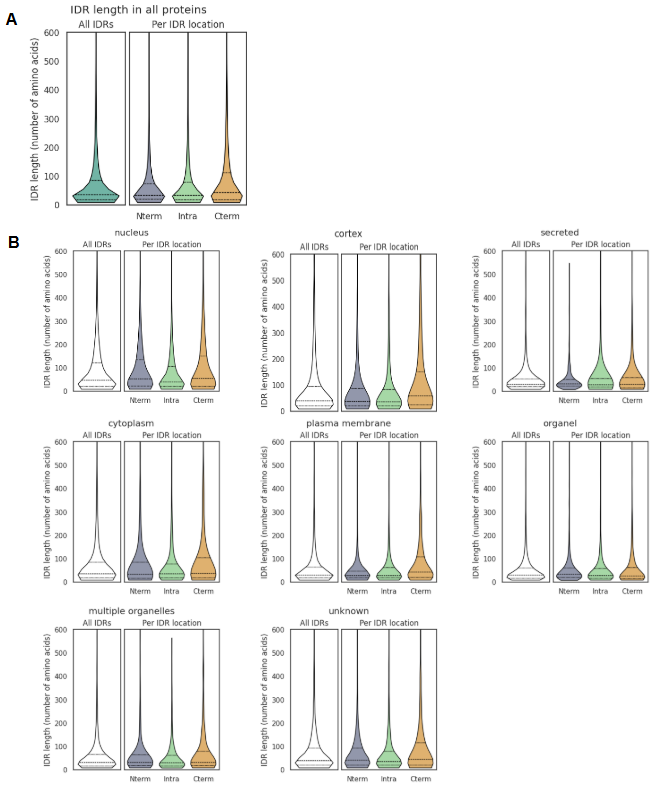

Supplemental Figure S4: Chemical properties of N-terminal (Nterm, purple), internal (Intra, green) and C-terminal (Cterm, orange) IDRs divided into cationic (R,H,K), anionic (D,E), polar (S,T,N,Q), hydrophobic (A,V,I,L,M,F,Y,W) and special (C,P,G) residues.

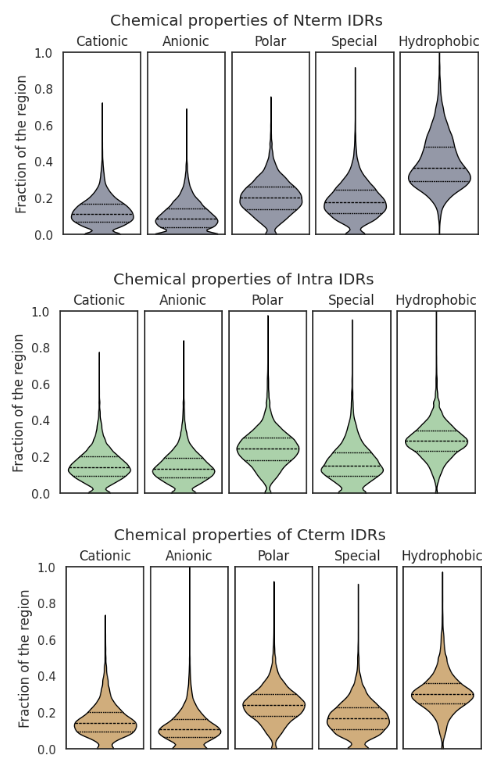

Supplemental Figure S5: AlphaMissense pathogenicity data, presented per amino acid type in either disordered regions (green) or ordered regions (grey). Areas under the curves are all set to 1. Consult Supplemental Table S3 for the relative occurrence of each amino acid type. The dark green and dark grey line indicate the distribution of all amino acids in disordered or ordered regions, respectively, and can be used to compare how the distribution of an individual amino acid is shifted relative to all amino acids in disordered or ordered residues.

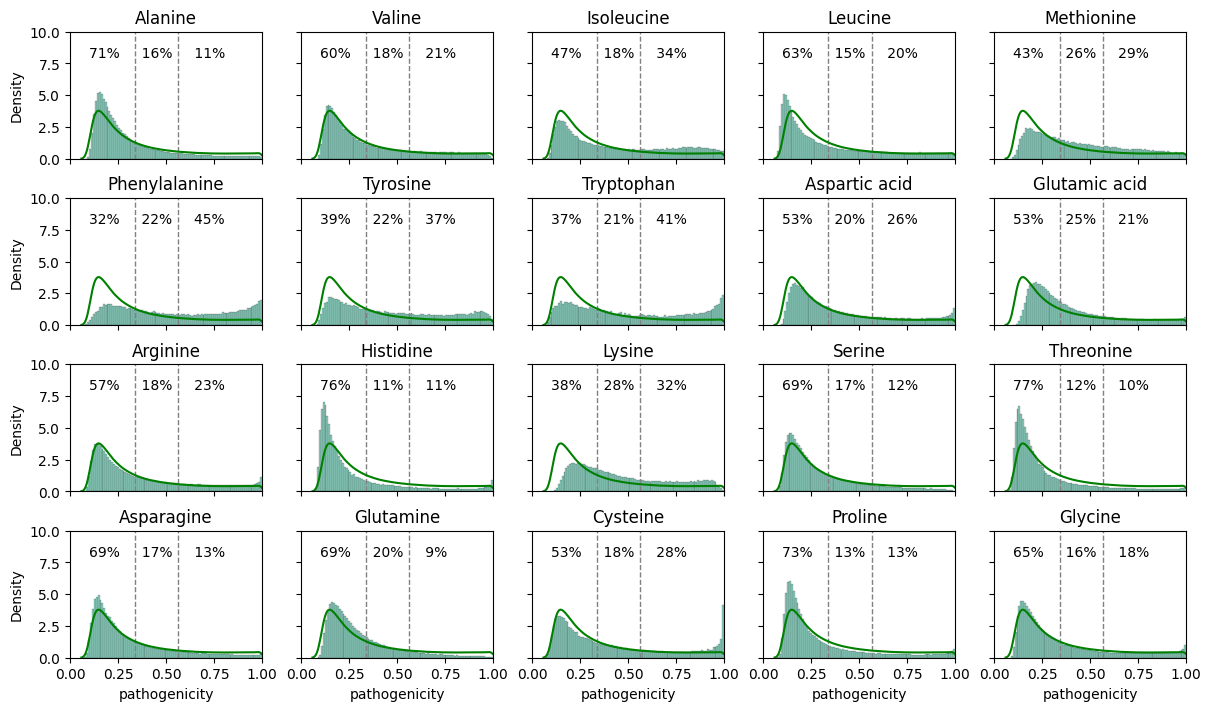

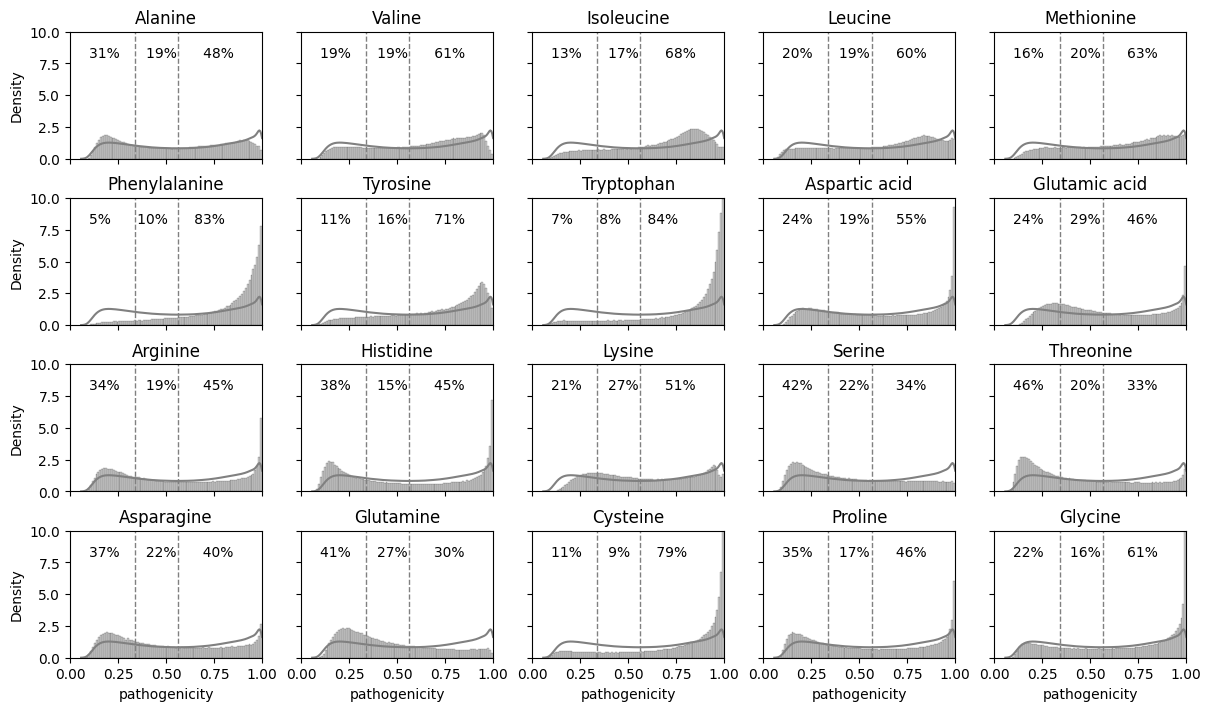

Supplemental Figure S6: PTMs found in IDRs (top row) and non-IDRs (bottom row) per subcellular localization of proteins. Note that phosphorylation is left out for clarity.

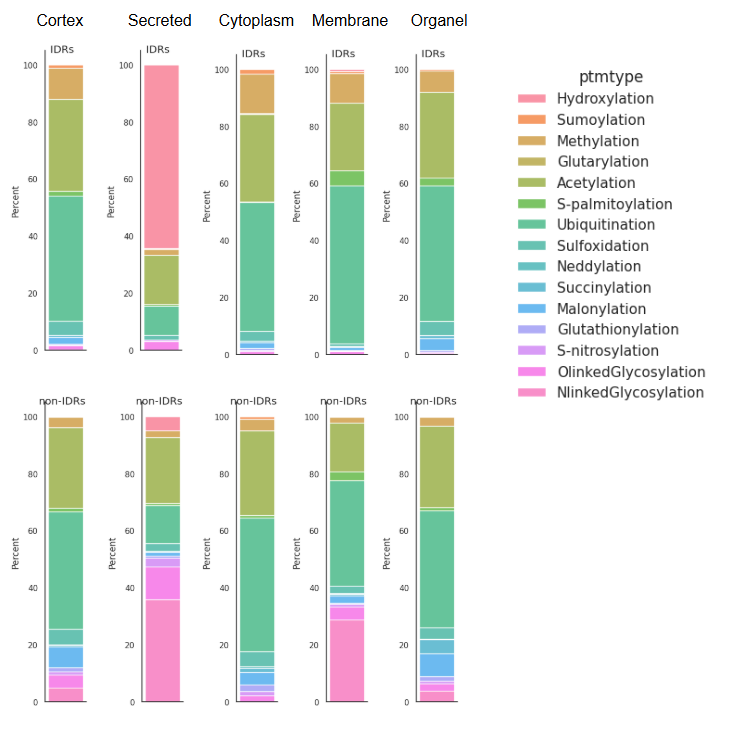

Supplemental Figure S7: Relative presence of phosphorylation in serine (S), threonine (T) or tyrosine (Y) in IDRs vs. non-IDRs. The dotted vertical lines mark the 50% border, and the numbers on the right-hand sides represent the number of PTMs found in total for that bar.

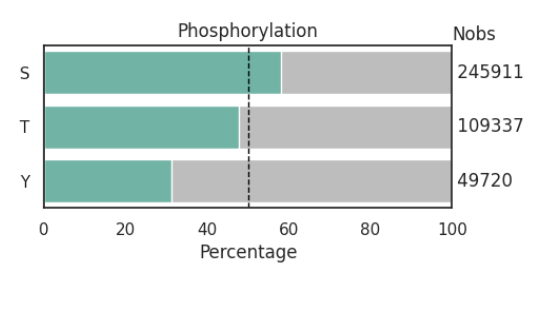

Supplemental Figure S8: Characteristics of proteins and IDRs selected in main Figure 5B. The IDRs selected (green top, right corner) are referred to as confidently ordered (c Ord) IDRs and the non-IDRs selected (grey, bottom left corner) are referred to as non-confident low-ordered (nC IOrd) non-IDRs. A) Cellular localization of c Ord IDRs (left) and nC lOrd non-IDRs (right). B) Length of all IDRs and non-IDRs, c Ord IDRs and nC lOrd non-IDRs, and their N-terminal (Nterm), internal (intra) and C-terminal (Cterm). C) Chemical properties of all IDRs and non-IDRs, c Ord IDRs and nC lOrd non-IDRs.

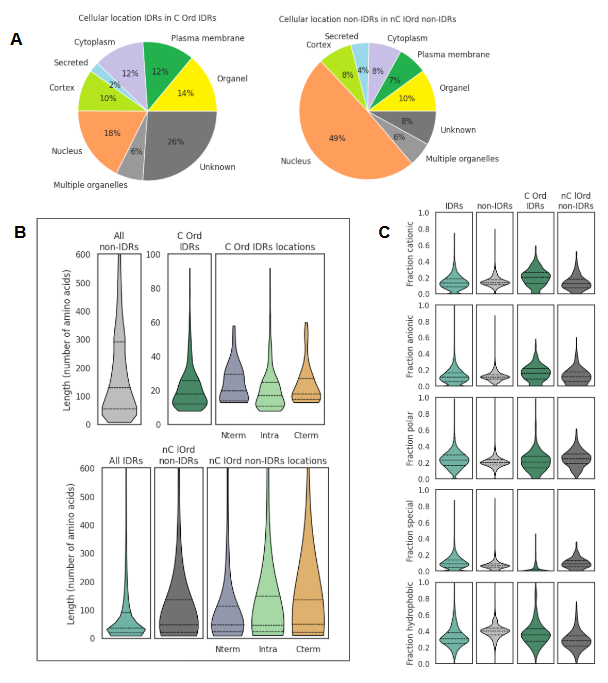

Supplemental Figure S9: A) Chemical properties of IDPs (pink) compared to the chemical properties of IDRs (green). From left to right the fraction of cationic, anionic, polar, hydrophobic and special residues found in the IDR or IDP is represented. B) Correlation between the percentage residues that adopt secondary structure (x-axis) and the average pLDDT score of the membrane IDPs (left) and IDPs with unknown location (right) found in our dataset. R^2^ marks the Pearson correlation coefficient.

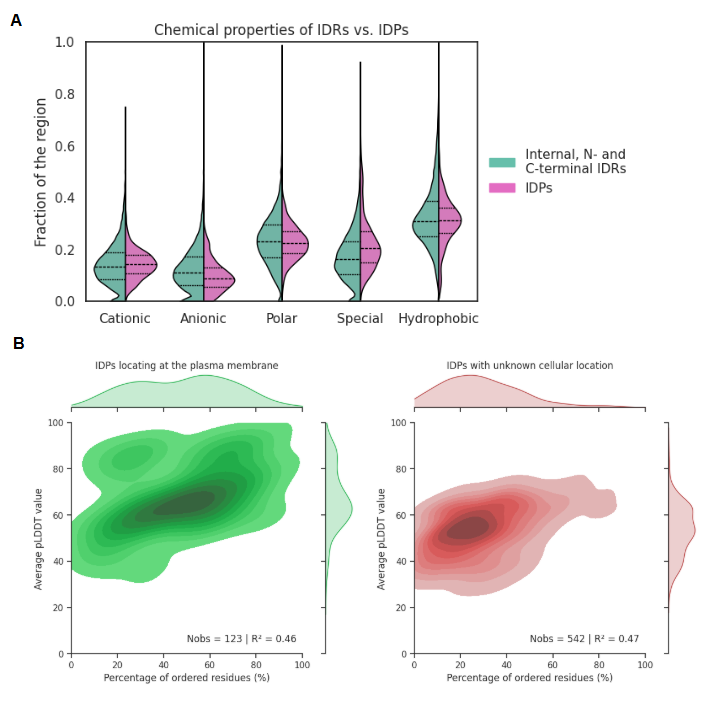

Supplemental Figure S10: A) Fraction of IDR residues in all IDR containing proteins (left) and in phase separation IDR containing proteins (right). B) Fraction of IDR residues in all IDR containing proteins (left) and in phase separation IDR containing proteins (right). C) Relative presence of methylation in IDRs vs. non-IDRs for all proteins and phase separating proteins. The dotted vertical lines mark the 50% border, and the numbers on the right-hand sides represent the number of PTMs found in total for that bar.

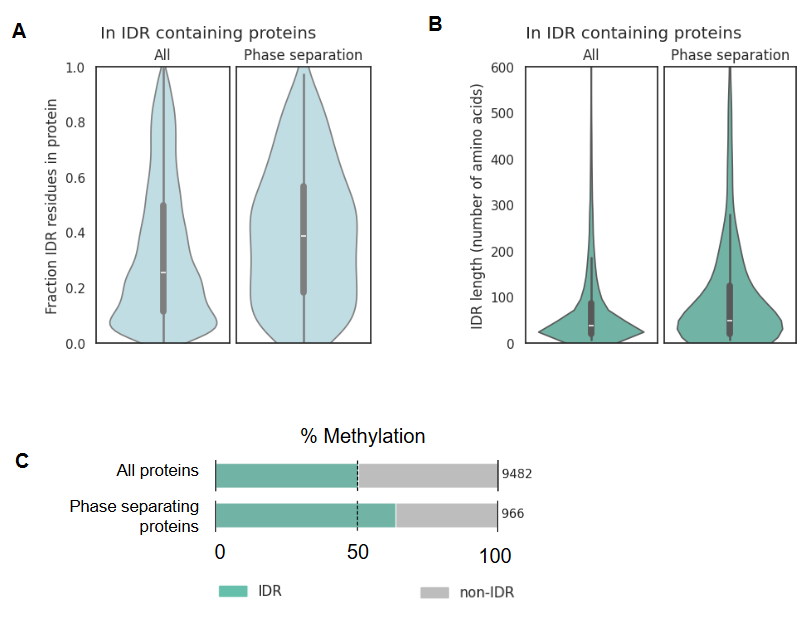

Supplemental Table S1: N-terminal IDRs that span >=90% of the protein

| Uniprot ID | Protein name | IDR fraction in protein | Number of amino acids in protein | IDR length |
| --- | --- | --- | --- | --- |
| D6RIA3 | Uncharacterized protein C4orf54 | 0.99 | 1793 | 1776 |
| O15231 | Zinc finger protein 185 | 0.91 | 689 | 629 |
| O95171 | Sciellin | 0.90 | 688 | 620 |
| P20382 | Pro-MCH | 0.91 | 165 | 150 |
| P35637 | RNA-binding protein FUS | 0.97 | 526 | 512 |
| Q8IXP5 | POU domain class 2-associating factor 2 | 0.95 | 288 | 274 |
| Q8TAM6 | Ermin | 0.92 | 284 | 262 |
| Q92681 | Regulatory solute carrier protein family 1 member 1 | 0.93 | 617 | 572 |
| Q96M34 | Testis-specific expressed protein 55 | 0.94 | 536 | 502 |

Supplemental Table S2: C-terminal IDRs that span >=90% of the protein

| Uniprot ID | Protein name | IDR fraction in protein | Number of amino acids in protein | IDR length |
| --- | --- | --- | --- | --- |
| A0A1B0GUW6 | Uncharacterized protein SPEM3 | 0.97 | 1196 | 1160 |
| C9JUS6 | Putative adrenomedullin-5-like protein | 0.91 | 153 | 139 |
| O00559 | Receptor-binding cancer antigen expressed on SiSo cells | 0.94 | 213 | 201 |
| P25063 | Signal transducer CD24 | 0.90 | 80 | 72 |
| Q05682 | Caldesmon | 0.97 | 793 | 768 |
| Q0VD83 | Apolipoprotein B receptor | 0.98 | 1097 | 1073 |
| Q13428 | Treacle protein | 0.96 | 1488 | 1432 |
| Q5D862 | Filaggrin-2 | 0.96 | 2391 | 2303 |
| Q5VYM1 | Spermatogenesis-associated protein 31G1 | 0.94 | 1079 | 1016 |
| Q6PRD7 | Cementoblastoma-derived protein 1 | 0.92 | 247 | 227 |
| Q7Z5B4 | Protein RIC-3 | 0.96 | 369 | 354 |
| Q8N350 | Voltage-dependent calcium channel beta subunit-associated regulatory protein | 0.92 | 705 | 647 |
| Q8N7X1 | RNA-binding motif protein | 0.92 | 1067 | 984 |
| Q8TAL5 | Uncharacterized protein C9orf43 | 0.92 | 461 | 423 |
| Q8TAV5 | Uncharacterized protein KCNJ5-AS1 | 0.95 | 145 | 138 |
| Q8TAX7 | Mucin-7 | 0.94 | 377 | 356 |
| Q99217 | Amelogenin | 0.96 | 191 | 183 |
| Q9BXS6 | Nucleolar and spindle-associated protein 1 | 0.90 | 441 | 397 |

Supplemental Table S3: Chemical properties of IDRs vs. non-IDRs per category (top) and per amino acid (bottom).

| Chemical property | Number of IDR amino acids | % of IDR amino acids | Number of non-IDR amino acids | % of non-IDR amino acids |
| --- | --- | --- | --- | --- |
| Polar | 817,863 | 24.78 | 1487,711 | 20.56 |
| Hydrophobic | 987,842 | 29.93 | 2917,578 | 40.32 |
| Special | 631,821 | 19.14 | 969,789 | 13,41 |
| Cationic | 458,176 | 13.88 | 1,024,230 | 14.15 |
| Anionic | 404,667 | 12.26 | 837,228 | 11.57 |
| Total | 3,30,0369 | 100 | 7,236,536 | 100 |

| Amino acid | Observations in IDRs | % in IDRs | Observations in non-IDRs | % in non-IDRs |
| --- | --- | --- | --- | --- |
| R | 195857 | 5.93 | 403433 | 5.57 |
| H | 77638 | 2.35 | 200540 | 2.77 |
| K | 184681 | 5.6 | 420257 | 5.81 |
| G | 257427 | 7.8 | 436025 | 6.03 |
| D | 152474 | 4.62 | 343622 | 4.75 |
| E | 252193 | 7.64 | 493606 | 6.82 |
| S | 363908 | 11.03 | 508867 | 7.03 |
| T | 179925 | 5.45 | 374973 | 5.18 |
| N | 104962 | 3.18 | 272021 | 3.76 |
| Q | 169068 | 5.12 | 331850 | 4.59 |
| A | 255789 | 7.75 | 488296 | 6.75 |
| V | 152300 | 4.61 | 472596 | 6.53 |
| I | 88366 | 2.68 | 367234 | 5.07 |
| L | 267355 | 8.1 | 787207 | 10.88 |
| M | 65445 | 1.98 | 161262 | 2.23 |
| F | 76576 | 2.32 | 311727 | 4.31 |
| Y | 53624 | 1.62 | 228760 | 3.16 |
| W | 28387 | 0.86 | 100496 | 1.39 |
| C | 54170 | 1.64 | 188103 | 2.6 |
| P | 320224 | 9.7 | 345629 | 4.78 |

Supplemental Table S4: Phase separating proteins without annotated IDRs.

| Uniprot ID | protname | Protein length | Essential |
| --- | --- | --- | --- |
| A6NCE7 | Microtubule-associated protein 1 light chain 3 beta 2 | 125 | No |
| O00444 | Serine/threonine-protein kinase PLK4 | 970 | Yes |
| O00487 | 26S proteasome non-ATPase regulatory subunit 14 | 310 | Yes |
| O14773 | Tripeptidyl-peptidase 1 | 563 | No |
| O43660 | Pleiotropic regulator 1 | 514 | Yes |
| P07320 | Gamma-crystallin D | 174 | No |
| P0DPB6 | DNA-directed RNA polymerases I and III subunit RPAC2 | 133 | No |
| P16402 | Histone H1.3 | 221 | No |
| P19484 | Transcription factor EB | 476 | No |
| P20226 | TATA-box-binding protein | 339 | Yes |
| P20671 | Histone H2A type 1-D | 130 | No |
| P28070 | Proteasome subunit beta type-4 | 264 | Yes |
| P29350 | Tyrosine-protein phosphatase non-receptor type 6 | 595 | No |
| P33240 | Cleavage stimulation factor subunit 2 | 577 | Yes |
| P35609 | Alpha-actinin-2 | 894 | No |
| P37198 | Nuclear pore glycoprotein p62 | 522 | Yes |
| P62491 | Ras-related protein Rab-11A | 216 | No |
| P63279 | SUMO-conjugating enzyme UBC9 | 158 | Yes |
| Q13618 | Cullin-3 | 768 | Yes |
| Q13702 | 43 kDa receptor-associated protein of the synapse | 412 | No |
| Q15370 | Elongin-B | 118 | Yes |
| Q6SZW1 | NAD(+) hydrolase SARM1 | 724 | No |
| Q8IXH7 | Negative elongation factor C/D | 590 | Yes |
| Q92879 | CUGBP Elav-like family member 1 | 486 | Yes |
| Q9GZQ8 | Microtubule-associated protein 1 light chain 3 beta | 125 | No |
| Q9UQM7 | Calcium/calmodulin-dependent protein kinase type II subunit alpha | 478 | No |
| Q9Y2X0 | Mediator of RNA polymerase II transcription subunit 16 | 877 | Yes |
| Q9Y572 | Receptor-interacting serine/threonine-protein kinase 3 | 518 | No |

Supplemental Methods S1: UniProt IDs that were deleted from the dataset for reasons described in the Methods section.

A0A087WUL8 A0A0A0MT78 A0A0A0MT89 A0A0A0MTA4 A0A0J9YX06 A0A0J9YXA8 A0A1B0GTB2 A0A3B3IS91 A2VEC9 A4UGR9 A6ZKI3 B1AH88 B7ZAP0 C0HLV8 F7VJQ1 L0R6Q1 L0R8F8 M0QZD8 O14686 O15018 O15050 O15230 O15417 O42043 O42043 O43149 O43687 O60229 O60281 O60494 O60613 O60673 O71037 O71037 O75445 O75592 O75691 O75962 O94915 O95071 O95278 O95359 O95467 O95613 O95714 O96033 P01266 P04114 P04275 P06881 P07203 P08519 P08F94 P0C7T4 P0DI83 P0DOY5 P0DP91 P0DPB5 P0DPF2 P0DPI4 P0DPQ6 P0DPR3 P10265 P11532 P12111 P13611 P15822 P15924 P18283 P20929 P20930 P21359 P21817 P22105 P22352 P24043 P25054 P25391 P35555 P35556 P36969 P42167 P42858 P46013 P46939 P49454 P49792 P49895 P49908 P50851 P51587 P55073 P58107 P58400 P58401 P59796 P59797 P60896 P61566 P61567 P61570 P61571 P61572 P61573 P61575 P61579 P61580 P61581 P61582 P61583 P62341 P62683 P62685 P62861 P63092 P63119 P63120 P63122 P63123 P63125 P63126 P63127 P63129 P63130 P63131 P63302 P78509 P78527 P78559 P84996 P87889 P98088 P98160 P98161 P98164 Q01484 Q02224 Q02388 Q02505 Q02817 Q03001 Q03164 Q07954 Q09666 Q0VDD8 Q12802 Q12830 Q12955 Q13315 Q13765 Q13948 Q14204 Q14315 Q14517 Q14571 Q14643 Q14789 Q15149 Q15413 Q15751 Q15772 Q15911 Q16787 Q16881 Q2LD37 Q3L8U1 Q4G0P3 Q4LDE5 Q5CZC0 Q5SZK8 Q5T011 Q5T1H1 Q5T4S7 Q5TBA9 Q5TFQ8 Q5THJ4 Q5VST9 Q5VT06 Q63HN8 Q685J3 Q68DQ2 Q69383 Q69384 Q6B8I1 Q6EEV4 Q6KC79 Q6N022 Q6V0I7 Q6V1P9 Q6ZNJ1 Q6ZQQ6 Q6ZR08 Q6ZRI0 Q6ZRS2 Q6ZS81 Q6ZTR5 Q709C8 Q70CQ2 Q70YC4 Q75N90 Q7LDI9 Q7Z407 Q7Z408 Q7Z5P9 Q7Z6Z7 Q7Z7G8 Q7Z7M0 Q86UP3 Q86UQ4 Q86VQ6 Q86WI1 Q86XX4 Q86YZ3 Q8IU53 Q8IVF2 Q8IVF4 Q8IWI9 Q8IYW2 Q8IZF6 Q8IZQ1 Q8IZQ5 Q8IZT6 Q8N2C7 Q8N2E6 Q8N3K9 Q8N726 Q8NCM8 Q8NDA2 Q8NEZ4 Q8NF91 Q8NFC6 Q8NFP9 Q8TCU4 Q8TD26 Q8TD57 Q8TDJ6 Q8TDW7 Q8TDX9 Q8TE73 Q8WUY3 Q8WWX9 Q8WXG9 Q8WXH0 Q8WXI7 Q8WXX0 Q8WZ42 Q902F8 Q92736 Q92813 Q96DT5 Q96E66 Q96JB1 Q96JG9 Q96JQ0 Q96L91 Q96M86 Q96N23 Q96PZ7 Q96Q15 Q96RL7 Q96RW7 Q96T58 Q99611 Q99698 Q99715 Q99996 Q9BQE4 Q9BQY6 Q9BVL4 Q9BXH1 Q9BXT5 Q9C0D9 Q9C0G6 Q9H251 Q9H496 Q9H5I5 Q9H799 Q9HC47 Q9HC84 Q9HCU4 Q9HDB5 Q9HDB8 Q9NNW7 Q9NR09 Q9NR48 Q9NR99 Q9NRC6 Q9NT68 Q9NU22 Q9NYC9 Q9NYQ6 Q9NYQ7 Q9NYQ8 Q9NZJ4 Q9NZR2 Q9NZV5 Q9NZV6 Q9P225 Q9P2D1 Q9P2D7 Q9P2P6 Q9UFH2 Q9UKH3 Q9UKN1 Q9UKN7 Q9UKZ4 Q9UMN6 Q9UPA5 Q9UPN3 Q9UQ35 Q9Y485 Q9Y493 Q9Y4A5 Q9Y4D8 Q9Y520 Q9Y6D0 Q9Y6I0 Q9Y6R7 Q9Y6V0 Q9YNA8 X6R8R1
